# Modeling Hsp70/Hsp40 interaction by multi-scale molecular simulations and co-evolutionary sequence analysis

**DOI:** 10.1101/067421

**Authors:** Duccio Malinverni, Alfredo Jost Lopez, Paolo De Los Rios, Gerhard Hummer, Alessandro Barducci

## Abstract

The interaction between the Heat Shock Proteins 70 and 40 is at the core of the ATPase regulation of the chaperone machinery that maintains protein homeostasis. However, the structural details of this fundamental interaction are still elusive and contrasting models have been proposed for the transient Hsp70/Hsp40 complexes. Here we combine molecular simulations based on both coarsegrained and atomistic models with co-evolutionary sequence analysis to shed light on this problem by focusing on the bacterial DnaK/DnaJ system. The integration of these complementary approaches resulted into a novel structural model that rationalizes previous experimental observations. We identify an evolutionary-conserved interaction surface formed by helix II of the DnaJ J-domain and a groove on lobe IIA of the DnaK nucleotide binding domain, involving the inter-domain linker.

## I. INTRODUCTION

The 70 kDa and 40 kDa Heat Shock Proteins (Hsp70/Hsp40) form the core of a chaperone machinery that plays essential roles in proteostasis and proteolytic pathways [1–3]. Hsp70s chaperones, and their cochaperone partners Hsp40s, are highly-conserved ubiquitous proteins, present in multiple paralogs in virtually all known organisms [1, 4]. The chaperoning role of this machinery is based on the ability of Hsp70s to bind client proteins in non-native states, thereby preventing and reverting aggregation, unfolding misfolded proteins, assisting protein degradation and translocation [5–8].

Members of the Hsp70 family are composed of two domains, connected by a flexible linker: The N-terminal nucleotide binding domain (NBD) binds and hydrolyzes ATP, whereas the C-terminal substrate binding domain (SBD) interacts with client proteins [9]. The nature of the bound nucleotide induces dramatically different conformations of Hsp70: In the ADP-bound state, the two domains are mostly detached and behave almost independently [10], whereas in the ATP-bound state, the SBD splits into two sub-domains that dock onto the NBD [11, 12]. Therefore, nucleotide hydrolysis and exchange result into large-scale conformational dynamics that regulates the chaperone interaction with client proteins [3]. Hsp40s are also called J-Domain Proteins, since they are invariantly characterized by the presence of a ~ 70 residue signature domain (J-domain), within a variable multi-domain architecture. This J-domain is composed of four helices (Fig. 1H). The two central helices II and III form an antiparallel bundle, connected by a flexible loop with a highly conserved distinctive histidine-proline-aspartate (HPD) motif [13]. Several studies have indicated the essential role of the J-domain and of the HPD motif in Hsp40/Hsp70 interactions [14–17]. While the structural diversity of Hsp40s mirrors the functional versatility of this complex machinery, the common conserved J-domain is strictly necessary for enhancing ATP hydrolysis by Hsp70 [4]. The modulation of Hsp70 ATPase activity through the formation of transient Hsp70/Hsp40/client complexes regulates the chaperone affinity for client proteins [18] and it is hence essential for all its multiple cellular functions.

**FIG. 1.**
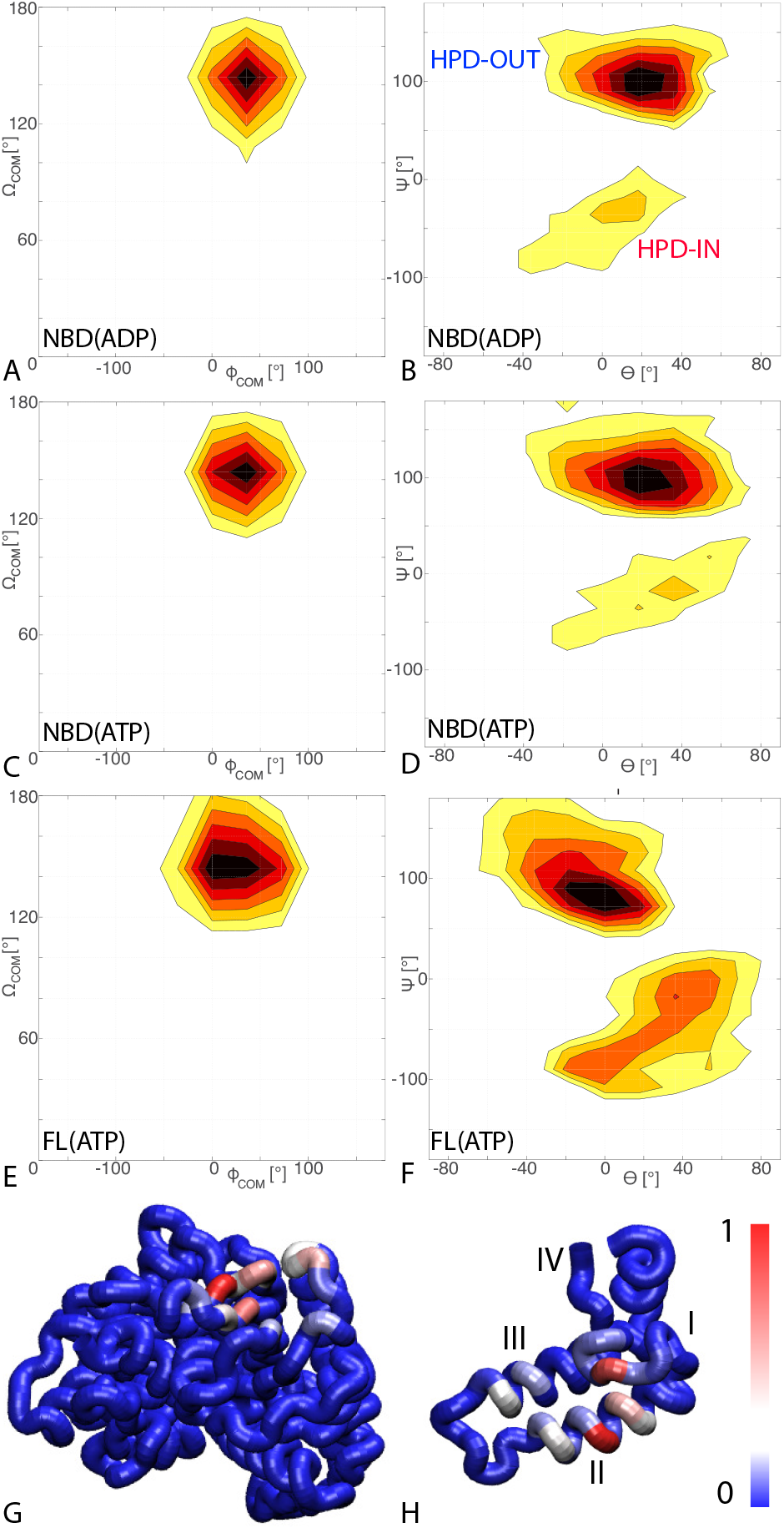
Binding modes of the JD/DnaK. A,C,E) The Free energy surfaces as a function of the spherical polar angles (Φ_*com*_, Ω_*com*_) of the JD center of mass for A) NBD(ADP), C) NBD(ATP), E) FL(ATP). The origin and the reference axes are defined by NBD center of mass and inertia axes, respectively. **B,D,F**) The Free energy surface as a function of Euler angles (Θ, Ψ) defining the relative orientation of J-domain w.r.t. the NBD for B) NBD(ADP), D) NBD(ATP), F) FL(ATP). The fixed and rotating coordinate systems are defined by the inertia axes of the NBD and JD respectively. Iso-lines are drawn at 1 (NBD(ADP/ATP)) or 1.2 (FL(ATP)) *k_B_T* intervals. See Fig. S4 for the third Euler angle. **G**) Probabilities of DnaK residues to be in contact with the JD (NBD(ATP) case). **H**) Probabilities of JD residues to be in contact with the DnaK (NBD(ATP) case). Helices I-IV of the JD are highlighted. See Fig. S2 for the NBD(ADP) and FL(ATP) cases.

Due to the continuous switching between multiple conformations [19], the chaperone-cochaperone interaction is intrinsically highly dynamic. Understanding the complex interplay between Hsp70 and Hsp40 at the mechanistic level is therefore a crucial task in order to gain a deeper functional understanding of this fundamental chaperone machinery.

Given the essential role of the Hsp70/40 interactions, extensive experimental evidence has been accumulated over the last two decades. Mutagenesis, surface plasmon resonance and NMR experiments have identified multiple putative interacting regions of the J-domain and Hsp70, mostly focusing on the E. Coli DnaK/DnaJ system [14, 16, 20–22]. In spite of this considerable effort, the dynamic and transient nature of the Hsp70/Hsp40 complex has posed severe challenges to their structural characterization and no consensus view has been reached yet.

To date, the only available high-resolution structure has been obtained by means of X-ray crystallography of the NBD of bovine Hsc70 (Hsp70) covalently-linked to the J-domain of bovine auxilin (Hsp40) [23]. However, this structure cannot be easily reconciled with NMR and mutagenesis data collected on DnaK/DnaJ, thus suggesting either major differences in the binding modes of bacterial and eukaryotic Hps70/40s or the trapping of a sparsely populated state that is influenced by non-native contacts [24, 25]. More recently, solution PRE-NMR experiments revealed another facet of the puzzle by identifying an alternative highly dynamic interface between ADP-bound DnaK and DnaJ [20].

Here we relied on both multi-scale molecular modeling and statistical analysis of protein sequences to shed light onto the details of the Hsp70/Hsp40 interactions. By combining these complementary techniques, we propose a structural model of the binding of bacterial DnaK/DnaJ, which is in good agreement with available experimental data and greatly extends our understanding of this elusive yet fundamental process.

## II. RESULTS

### A. Coarse-grained simulations identify binding regions and suggest structural models for the DnaK/J complex

We firstly characterized DnaK/DnaJ interactions by means of Monte Carlo simulations based on a Coarse Grained (CG) model combined with effective energy potentials validated against structural and thermodynamic properties of protein complexes with low binding affinity [26–28]. In this scheme, binding partners are modeled as rigid bodies, using one interaction site per amino-acid (residue) located at the *C_α_* position of the experimental structure. Intermolecular energy functions are based on statistical contact potentials and long-range Debye-Hückel electrostatic interactions [26]. A replica exchange Monte Carlo simulation protocol is adopted to exhaustively sample all the relevant bound conformations. Here, we took advantage of this approach and of the availability of high-resolution structures to investigate complexes formed by the J-domain of E. Coli DnaJ (JD) with the DnaK NBD, both in its ADP- and ATP-bound conformations (NBD(ADP), NBD(ATP)). Moreover, we extended this analysis to full-length ATP-bound DnaK (FL(ATP)) in order to unveil a possible role of the SBD in the binding process.

CG trajectories were analyzed to determine the binding affinity of the three DnaK constructs for JD and to characterize the most favorable complex conformations. Calculated binding affinities (Kd=540μM ± 60 NBD(ADP), Kd=370μM ± 35 NBD(ATP), Kd=23μM ± 3 FL(ATP)) are compatible with previous experimental determinations [14, 20, 29] and their significant dependence on the presence of the SBD and linker suggests a stabilizing role of this region in the DnaK/JD complex. From the structural point of view, the analysis of the conformational ensembles corresponding to bound complexes revealed several distinctive features of the DnaK/JD binding process, along with a certain degree of conformational heterogeneity (Fig. S1). Notably, the free-energy surfaces as a function of NBD-centered spherical coordinates (Fig.1A,C,E) clearly indicate that a specific binding site predominates in all the simulated DnaK constructs, irrespectively of the bound nucleotide and of the presence of the SBD and linker.

In order to better characterize this favoured binding interface, we calculated the probability of each DnaK residue to be in direct contact with JD and mapped it onto the NBD structure (Fig. 1G). These results suggest that the formation of DnaK/DnaJ complexes mostly involves a DnaK region located on lobe IIA of the NBD, in a negatively charged narrow groove formed by a beta-sheet and a short loop (Fig. 1G, 2A,B). Furthermore, analysis of the complementary interface on JD suggests that its interaction with DnaK is mostly mediated by the positively-charged helix II and few residues on helix I (Fig. 1H).

The conformational ensembles obtained by CG simulations were further analyzed to identify the most relevant orientations of the JD in the bound complexes. Particularly, the free-energy surfaces as a function of Euler angles measuring the relative orientation of the binding partners reveal two major conformational subensembles in all the simulated systems (Fig. 1B,D,F, Fig. S4). Cluster analysis of the CG trajectories suggests that these distinct intermolecular arrangements correspond to complexes with similar binding interfaces but opposite orientations of the JD with respect to DnaK (Fig. 2, Fig. S3). Indeed, the conserved HPD loop of the JD points outwards in one conformational sub-ensemble (HPD-OUT, Fig. 2A,C) whereas in the other it is close to the groove on NBD where the interdomain linker docks (HPD-IN, Fig. 2B,D). These two arrangements were observed in all the simulated systems with very limited perturbations due to the presence of SBD and they together account for more than 91% of the populations in the bound ensembles. In all the systems, the population of the HPD-OUT conformation is higher than that of HPD-IN. However, the limited free-energy differences between the two binding modes (1.5 kcal/mol) are comparable to the expected uncertainty of the CG model [26] and thus do not allow us to draw definitive conclusions about their relative stability.

**FIG. 2.**
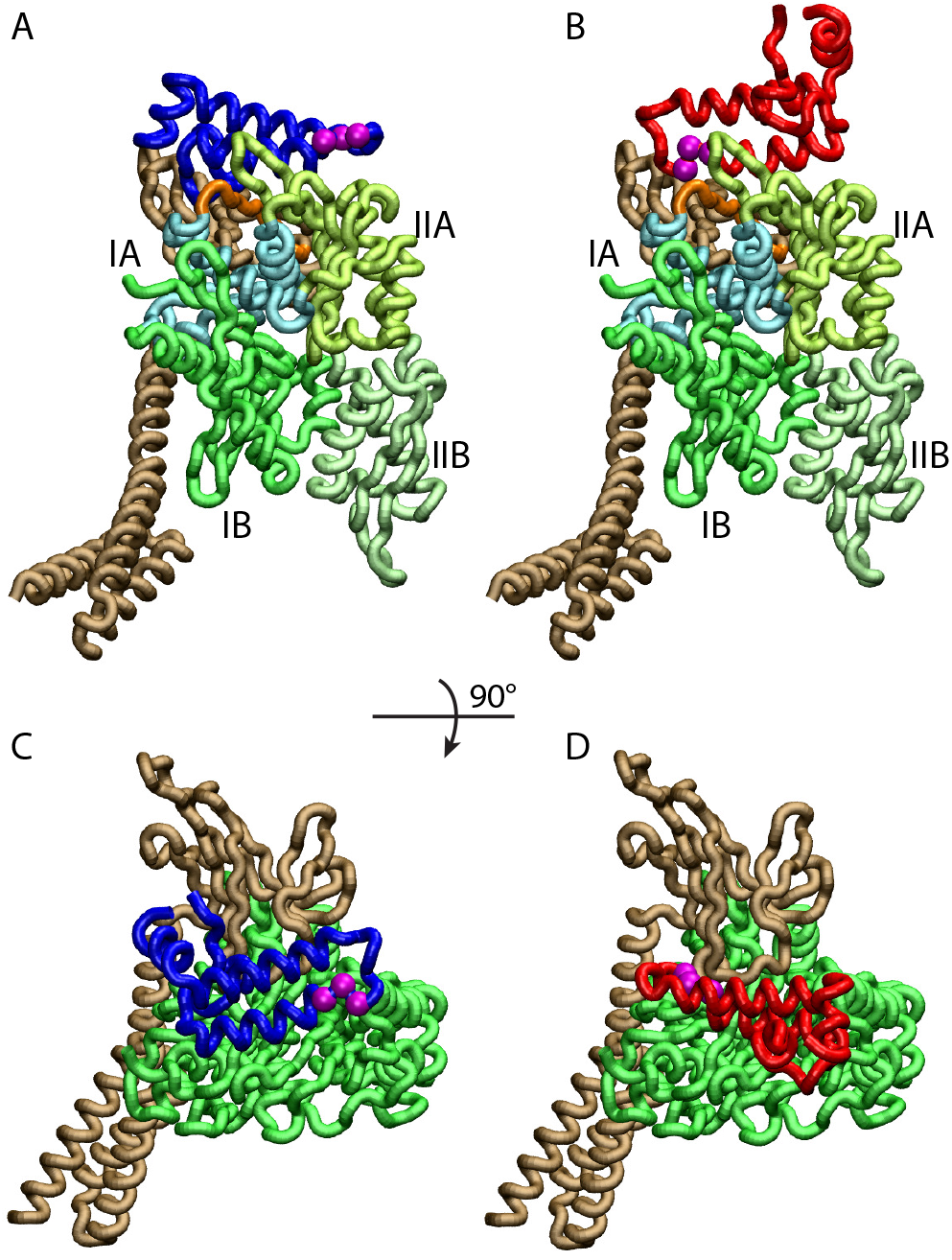
Conformations of the DnaK:JD complex. Most representative HPD-OUT/IN conformations for the FL(ATP):JD system (HPD-OUT: A) and C), HPD-IN: B) and D)). The four lobes forming the sub-structures of the NBD are highlighted. The SBD is in brown. The docked inter-domain linker is in orange. The HPD tripeptide of the J-domain is depicted as magenta spheres. The JD is depicted in blue (HPD-OUT) or red (HPD-IN). See Fig. S3 for the NBD(ADP) and NBD(ATP) systems. For readability, the NBD in the rotated panels C) and D) is only colored in green.

### B. Coevolutionary Analysis predicts conserved DnaK-DnaJ contacts

Statistical analysis of the covariation in multiple sequence alignments (MSAs) represents an extremely valuable approach to investigate protein structure by identifying residue-residue interactions that are evolutionarily conserved [30, 31]. Particularly, Direct Coupling Analysis (DCA) [32, 33] of paired MSAs of interacting proteins has been successfully applied to predict interfaces of protein complexes [34, 35]. The canonical matching algorithms for generating paired MSAs are based on intergenic distances and they unfortunately cannot be directly applied to Hsp70/40 interaction, due to the promiscuity and the general lack of operon structure in this family. In order to circumvent this difficulty, we adopted an alternative approach based on the generation of an ensemble of stochastically-matched MSAs. In this context, the statistical reliability of inter-residue couplings can be related to their frequency of appearance within the DCA predictions obtained from all the realizations of the MSA ensemble (see Methods for details).

Here we took advantage of the large sizes of the Hsp70/40 families (Hsp70: 20061 sequences, Hsp40: 26254 sequences, see Methods) to evaluate inter-residue evolutionary couplings between the Hsp40 JD and the Hsp70 NBD. The high degree of conservation of these domains guarantees accurate alignments and thus reliable results. This analysis identified three inter-protein residue pairs that stand out among coevolving pairs in the Hsp40 and Hsp70 families (Fig. 3A), corresponding to N187-K23, D208-K26 and T189-R19 in E.Coli DnaK and DnaJ numbering. The spatial proximity of N187/D208/T189 on the DnaK NBD, as well as the proximity between K23/K26/R19 on the JD, suggest the presence of well-defined binding patches that are evolutionary-conserved across Hsp40/70 families.

**FIG. 3.**
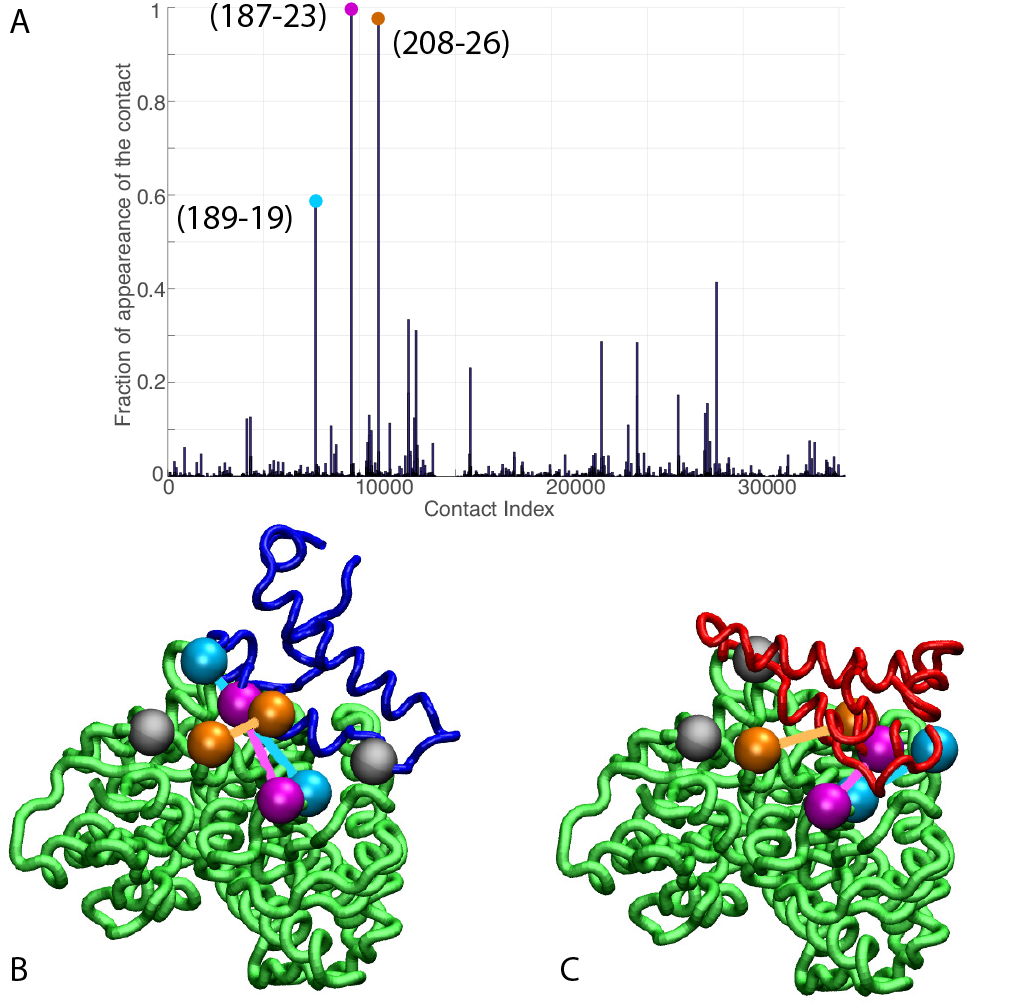
Coevolutionary predicted contacts. **A)** Frequency of appearance of the coevolutionary inter-protein contacts (see Methods). The three most frequent contacts are highlighted (N187-K23 magenta, D208-K26 orange and T189-R19 blue. Numbering refers to the E.Coli DnaK-DnaJ (Uniprot IDs: DnaK P0A6Y8, DnaJ P08622)). **B-C)** The same three contacts represented on the HPD-OUT (blue, panel B) and HPD-IN (red, panel C) conformations of the NBD(ATP):JD complex. Coevolving residues are depicted by spheres, following the color scheme of panel A). Grey spheres represent Asp35 of the 33HPD35 motif on JD and Arg167 of the NBD.

Even more remarkably, these patches are perfectly overlapping with the binding regions predicted by CG modeling, i.e., helix II in DnaJ and a sub-region of lobe IIA in DnaK NBD. Thanks to this overall excellent agreement, DCA predictions can be used to evaluate the HPD-IN and HPD-OUT binding modes suggested by CG simulations (Fig. 3B,C). Quantitative assessment is limited here by several factors such as the difficulty of translating coevolutionary couplings into exact distance restraints, the limited resolution of the residue-based CG model and the dynamic nature of the JD/DnaK complexes. Nevertheless, while both the intermolecular conformations might be compatible with DCA predictions, the specific binding pattern predicted by coevolutionary analysis matches significantly better the JD orientation observed in HPD-IN (Fig. 3C). This conclusion can be further strengthened by the observation that the average *C_α_ – C_α_* distance observed in HPD-OUT for the pair T189-R19 seems too large (> 20Å) to justify a direct interaction, and thus a strong statistical coupling between those residues.

### C. Atomistic MD validate the stability of the docked conformations

To further assess the reliability of the predicted docked-conformations, we performed atomistic Molecular Dynamics (MD) simulations of the DnaK-DnaJ complexes. Atomistic structures corresponding to the most relevant bound conformations observed in CG simulations were obtained using the RosettaDock algorithm (See Methods). Particularly, for each system (JD:NBD(ADP), JD:NBD(ATP) and JD:FL(ATP)) we generated sets of 10 atomistic structures for both HPD-IN and HPD-OUT binding modes in order to best represent the conformational fluctuations observed at the CG level. We then used these representative conformers to generate all-atom explicit-solvent MD trajectories of 30 ns (aggregated simulation time of 1.8 μs). Even if this limited time scale clearly prevents an exhaustive characterization of the inter- and intramolecular conformational dynamics, the multiple MD runs provide valuable information about the differential stability of the various complexes.

Effectively, trajectories of the NBD(ADP):JD and NBD(ATP):JD complexes starting from the HPD-OUT conformations display high conformational variability (Fig. 4A,C) whereas those started from HPD-IN conformations are significantly more stable in the simulated time scale (Fig. 4 B,D). This difference is absent or significantly less pronounced in the simulations of full-length DnaK (Fig. 4E,F) likely due to the stabilizing effect of JD-SBD interactions. These qualitative indications were corroborated by structural analysis of the last part (10ns) of the MD trajectories (Tab. I, Fig. S5, Fig. S6). Notably, average *C_α_* Distance Root Mean Square deviation (dRMS) validated the previously mentioned trend and revealed in all the systems a non-negligible amount of conformational dynamics (Tab. I, Fig. S5). To better quantify the degree of intramolecular dynamics we measured the average angular deviation Θ of the JD with respect to its orientation in the corresponding reference CG conformation (Tab. I, Fig. S6). This analysis further confirms that the HPD-IN docking mode is stable in all the considered complexes whereas the HPD-OUT intermolecular conformation is significantly distorted in absence of the SBD.

**FIG. 4.**
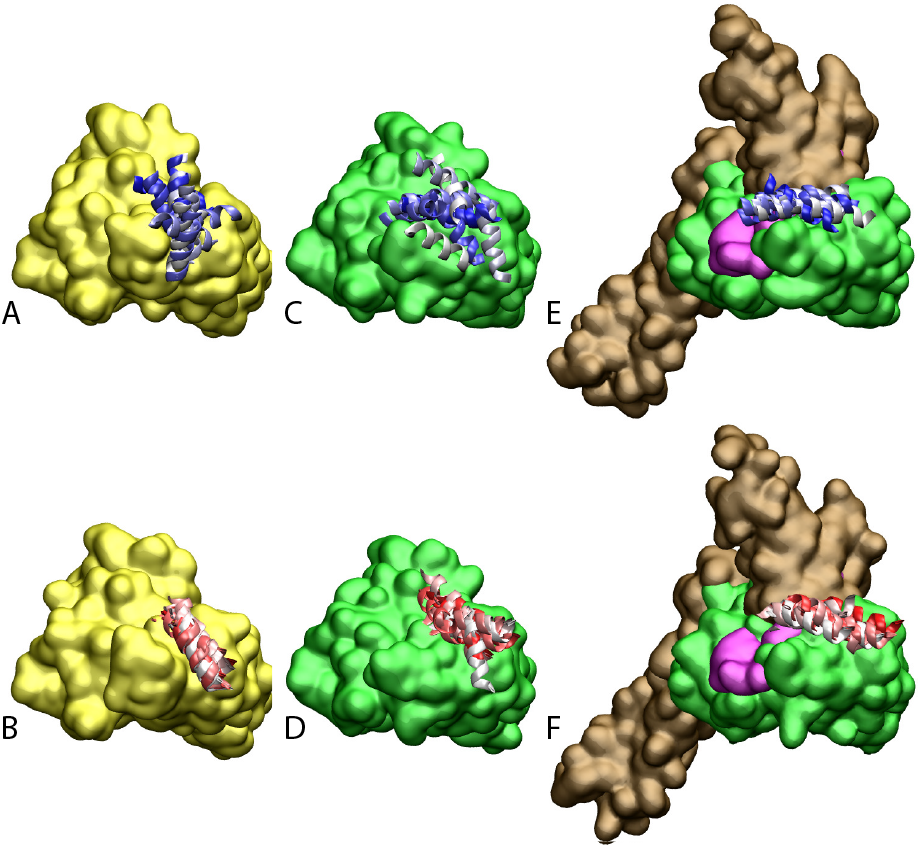
Atomistic stability analysis. Final frames of the 10 all-atom MD trajectories. **A**) HPD-OUT/NBD(ADP) **B**) HPD-IN/NBD(ADP) **C**) HPD-OUT/NBD(ATP). d) HPD-IN/NBD(ATP). e) HPD-OUT/FL(ATP). **F**) HPD-IN/FL(ATP). For ease of readability, only helices II and III of the JD are depicted. Rainbow coloring (white to blue for HPD-OUT, white to red for HPD-IN) is used to better differentiate the 10 frames. The DnaK NBD is colored in yellow for NBD(ADP) and in green for NBD(ATP) and FL(ATP). The docked inter-domain linker is colored in magenta in FL(ATP). The SBD is colored brown in FL(ATP).

**TABLE I.** Stability analysis of atomistic MD. dRMS denote the distance Root Mean Square deviation between *C_α_* atoms of the JD and DnaK. Θ denotes the angle formed by the principal axis of the JD with respect to the principal axis of the JD in the reference CG conformation. Standard deviations over the 10 MD trajectories are reported in parenthesis. See SI methods for details.

| | dRMS [$\text{\AA}$] | $\Theta$ [$^\circ$] |
| --- | --- | --- |
| NBD(ADP) HPD-OUT | 9.2 (2.1) | 40.7 (17.9) |
| HPD-IN | 3.7 (0.9) | 16.8 (4.4) |
| NBD(ATP) HPD-OUT | 12.3 (5.0) | 43.7 (12.8) |
| HPD-IN | 6.1 (3.0) | 19.3 (13.4) |
| FL(ATP) HPD-OUT | 6.2 (2.7) | 18.5 (9.2) |
| HPD-IN | 5.2 (2.0) | 17.6 (10.4) |

## III. DISCUSSION

The integration of complementary approaches such as co-evolutionary sequence analysis and molecular modeling both at the coarse-grained and atomistic scale allowed us to shed light on the structural details of the crucial, yet elusive, interaction of DnaJ with DnaK. Indeed, molecular simulations based on a CG model specifically suited to study low-affinity protein binding identified the positively charged helix II of the JD and a region close to lobe IIA of the DnaK NBD as the most relevant interaction sites in the formation of DnaK/JD complexes. This prediction was further confirmed by DCA results showing that at least three residue-residue interactions across this interface are conserved by evolution in the Hsp40/70 families. These findings are in good agreement with several experimental evidences collected in the last twenty years. Indeed, a major role for helix II of JD in the DnaK/DnaJ interaction has been suggested both by NMR and mutagenesis experiments [14, 16]. Furthermore, recent PRE-NMR investigation of the interaction of JD with ADP-bound DnaK indicated the region ^206^EIDEVDGEKTFEVLAT^221^ as the main binding region on DnaK [20]. While our prediction is in excellent agreement with this evidence, the observation that the same interaction site was present in all the simulated systems (ATP- and ADP-NBD and ATP-bound full-length DnaK) strongly suggest that this region is likely to play a primary role throughout the chaperones functional cycle, thus greatly extending its physiological relevance. Interestingly, the predicted bound conformations located the J-domain in near proximity of the docked inter-domain linker in FL(ATP) (Fig. 2), which has been shown to play a central role in the allosteric coupling of the two domains in the Hsp70s cycle [36–38]. Either direct interactions between the inter-domain linker and the the JD, or their competitive binding to the same region of Hsp70 NBD, are compatible with a regulatory role of the JD in the allosteric cycle of Hsp70s. The dynamical interplay between NBD, inter-domain linker and JD, and their mechanistic roles in the exact regulation of ATP hydrolysis and client binding remains however unclear, and calls for extended numerical and experimental investigations.

Beyond a detailed characterization of the binding regions on DnaK and the JD, our integrated approach provided precious information about the interprotein arrangement in the transient DnaK/JD complexes. Effectively, CG modeling suggested two possible binding modes characterized by opposite orientations of the JD (Fig. 2). Both these putative conformations were only minimally affected by structural differences in the NBD upon ATP/ADP binding or by the interdomain docking in full-length ATP-bound DnaK. Direct comparison of these results with the interaction pattern inferred from coevolutionary analysis reveals an excellent agreement for one of the conformations (HPD-IN). This first indication was further verified by performing MD simulations based on accurate atomistic potentials and explicit treatment of the solvent. Although the computational cost of this approach limits its predictive capabilities in the study of protein-protein interactions, the analysis of multiple MD trajectories showed a high structural stability of the HPD-IN conformations. Further elements supporting the relevance of this structure can be found by taking into account the role of the highly-conserved HPD loop of the JD in the dynamical chaperone/cochaperone interactions. Indeed, several mutagenesis studies have shown that the HPD loop is fundamental for functional chaperone/cochaperone interactions [16, 39]. NMR investigations have reported conflicting evidences about the actual involvement of the HPD region in the Hsp70/Hsp40 interface [14, 20, 40]. However, the observation that the DnaK R167N mutation could suppress the deleterious effect of the DnaJ D35N mutation strongly pointed toward a direct, yet possibly transient interaction of these residues during the chaperone functional cycle. Strikingly, this finding is perfectly compatible with the spatial proximity of DnaJ D35 and DnaK R167 in HPD-IN conformation (Fig. 3C) whereas it cannot be easily reconciled with the orientation of the HPD observed either in the HPD-OUT binding mode (Fig. 3B) or in the average solution structure of the DnaK:JD complex proposed on the basis of PRE-NMR experiments [20]. The HPD-IN conformation can hence best recapitulate the most relevant experimental evidences about prokaryotic Hsp70/Hsp40 systems and provides a suggestive model for the elusive DnaK/DnaJ complex.

The alternative arrangement observed in the bovine auxilin:Hsc70 complex [23], and its poor agreement with NMR/mutagenesis data on DnaK/DnaJ [14, 20] raise the question of the uniqueness of the Hsp70/Hsp40 binding mode [41]. Whether these major structural differences are due to artifacts introduced by the artificial cross-linking, to the existence of multiple dynamic interaction interfaces, or to phylogenetic differentiation of Hsp70/40s remains an essential yet unsolved question. In this respect, the successful combination of coevolutionary and molecular modeling analysis proposed here paves the road for further analysis to tackle these challenges.

### A. Materials and Methods

#### 1. Coarse-Grained Simulations

We used the coarse-grained model introduced in [26] to simulate the binding of DnaJ to DnaK constructs. Both proteins were treated as rigid bodies, at a resolution of one bead per residue centered on the *C_α_* atoms. We modeled NBD(ADP) using the structured region (residues 4–380) of ADP-bound DnaK (pdb: 2kho [10]) whereas we relied on the X-ray structure of ATP-bound DnaK (pdb: 4jne [12]) for both NBD(ATP) (residues 4–380) and FL(ATP) (residues 1–600).

Conformations were sampled from the equilibrium distribution using a replica-exchange Monte-Carlo (REMC) algorithm in a prediodic box, with 20 replicas distributed in the temperature range 200–395K. A total of 2·10^6^ MC-steps were performed for each replica and samples were recorded every 100 steps. Dissociation constants were calculated by measuring the fraction of bound conformations, and simulations were repeated with five increasing box sizes (240–360 Å for NBD, 300–420 Å for FL(ATP)). Bound conformations were extracted by selecting all complexes in which the two proteins had at least one pair of beads within 8 Å distance and total interaction energy equal or below −2*k_B_T*. All subsequent analysis on the CG complexes have been performed on the ensemble of bound complexes. The GROMOS algorithm [42] with a cutoff radius of 5 Å was used to perform cluster analysis of the CG trajectories. See SI for extended details of the coarse-grained simulations.

#### 2. Coevolutionary Analysis

Sequences for both Hsp40 and Hsp70 were retrieved from the Uniprot/Swissprot databases using the HMMER package [43] with manually curated seeds. After filtering of gapped sequences, the multiple sequence alignments (MSAs) contained 20061 (Hsp70) and 26254 (Hsp40) sequences.

To build a stochastically matched MSA, we used the following protocol: For each organism, a random sequence of Hsp70 was selected and randomly matched to a single Hsp40 sequence of the same organism. These two sequences were then removed from the pool of available sequences in the current organism. This procedure was then repeated until there were no more available Hsp40 or Hsp70 sequences to match in the current organism. We then repeated this random matching for all organisms possessing at least one Hsp40 and Hsp70 sequence. Direct-Coupling Analysis (DCA) was performed on each of the 1000 stochastically concatenated MSAs using the asymmetric version of the pseudo-likelihood method [44], with standard parameters (maximum 90% sequence identity, regularization parameters λ_*H*_ = λ_*J*_ = 0.01). For each realization, significant predicted contacts were selected using the inter-protein selection criterion introduced in [34], with a threshold set at 0.8.

The selection frequencies of all inter-protein contacts were computed over the 1000 realizations (Fig. 3C) and the most frequently appearing contacts were selected for further analysis. To set a threshold on the number of selected top appearing contacts, we computed the solvent accessible surface area (SASA) of the pairs of residues involved in the top appearing contacts. We then selected all ranked contacts before the appearance of a buried residue (SASA < Å^2^) in the contact pair. This resulted in the selection of 3 significantly conserved DCA predicted contacts (Fig. 3A,B,C). See SI for details on the coevolutionary analysis.

#### 3. Atomistic Simulations

For all the simulated systems, we used the RosettaDock [45, 46] protocol to obtain atomistic structures from the low resolution CG conformations corresponding to the HPD-IN and HPD-OUT binding modes. Particularly, we took advantage of the multiscale docking protocol [45] to generate 1000 all-atom conformation from the CG structures corresponding to the center of each HPD-IN and HPD-OUT cluster. We then selected the 10 best scoring structures among those within a deviation equivalent to the radius of the clusters (*C_α_* RMSD ≤ 5Å) and we solvated them in dodecahedral boxes containing approximately 26000 and 60000 water molecules for NBD:JD and FL:JD complexes, respectively. MD simulations were performed using the GROMACS 5 MD package [47], with the AMBER14 force-field [48] and TIP3P water model [49]. Given the large internal dynamics of Hsp70, we used harmonic restraints on the backbone atoms of the Hsp70 constructs (NBD(ADP),NBD(ATP) and FL(ATP)) in order to focus on the inter-protein dynamics. Additional details of the MD parameters and system equilibration are provided in SI.

## Supplemental Information

### I. COEVOLUTIONARY ANALYSIS

#### A. Sequence Extraction and Preprocessing

To perform Direct-Coupling analysis we used the following sequence extraction protocol: We built two separate seeds containing Hsp70 (resp. Hsp40) sequences, covering a broad portion of the tree of life. Hidden Markov Models (HMM) were then built for each protein family using the HMMER package [43]. The union of the Uniprot and Swissprot databases were then scanned for homologues using these HMMs. This resulted in multiple-sequence-alignments for the Hsp70 and Hsp40 families containing respectively 20061 and 26254 homologues. Both MSAs were then further filtered, removing all sequences containing more than 10% of gaps. Finally the Hsp70 MSAs was cropped, keeping only the positions corresponding to the nucleotide-binding domain and linker. The resulting MSAs covered the following ranges of the E.Coli DnaK/DnaJ proteins: DnaK (Uniprot ID: P0A6Y8) I4-T395, DnaJ (Uniprot ID: P08622) K3-G78.

#### B. Random Paralog matching

To detect inter-protein coevolving residue pairs, concatenated MSAs of interacting protein sequence pairs must be built. Given the absence of known interaction network of Hsp40s and Hsp70s and the lack of conservation of the number of paralogs throughout species, no trivial matching could be performed. Furthermore, the approach of matching interacting sequence pairs based on their genomic proximity [34, 35, 50] failed due to the lack of operon organization of Hsp70s and Hsp40s. We therefore employed a stochastic approach to the sequence-matching problem, which consists of the following steps:

- For each organism:
  Randomly select a sequence of Hsp70 and randomly matched it to a single Hsp40 sequence of the same organism.
  Remove these two sequences from the pool of available sequences in the current organism.
  Repeat this procedure until there are no more Hsp40 or Hsp70 sequences to match in the current organism.
  Repeat the procedure for all organisms possessing at least one Hsp40 and Hsp70 sequence.

This procedure generated a stochastic realization of a matched MSA, ensuring that each sequence was present only once in the MSA. This constraint of matching each sequence only once avoided the combinatorial explosion of the size of the random MSAs and consequently the dilution of the coevolutionary signal due to the presence of an overwhelming majority of non-interacting protein pairs. We generated 1000 such random MSAs and performed DCA on each of them individually. Significantly coevolving residue pairs were then selected using the inter-protein contact selection criterion introduced in [34], with cutoff set at 0.8. We then computed the selection frequency for each contact and retained the residue-pairs most often selected for subsequent analysis. The rationale behind this procedure was that contacts appearing repeatedly in multiple random realizations were robust to matching noise in the MSAs and should therefore reflect a strong coevolutionary underlying signal.

### II. ATOMISTIC SIMULATIONS

#### A. Simulation protocol

All simulations have been performed in a dodecahedral box with periodic boundary conditions. Simulations were carried out with the following protocol:

- Starting structures were solvated with TIP3P water molecules and subsequently energy minimized by steepest descent.
- A first NVT equilibration phase (1ns) was performed, putting full restraints on all proteins, ATP (when present) and MG atoms.
- A second NPT equilibration phase (1ns) was performed keeping the same restraints as in the 1ns NVT equilibration phase.
- Subsequently, another NPT equilibration was performed (10ns), putting restraints on the protein backbone only (DnaK and DnaJ).
- Finally, production runs were carried out for 30ns, keeping only restraints on the DnaK backbone.

Temperature was kept constant (T = 300K) using the v-rescale thermostat and NPT (P=1atm) simulations relied on a Parrinello-Rahman thermostat [51]. The equations of motion were integrated with a time step of 2 fs. All covalent bonds were constrained to their equilibrium values using the LINCS algorithm [52]. The electrostatic interactions were calculated by the Particle Mesh Ewald algorithm, and a cutoff of 10 nm was used both for LennardJones interaction and for the real-space coulomb contribution.

The Distance Root Mean Square (dRMS) measurements were calculated by

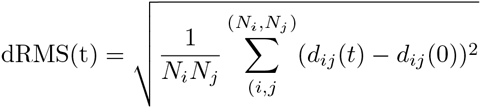

where *i* (resp. *j*) are indexes of the residues belonging to the J-domain (resp. DnaK), and *d_ij_(t)* denotes the distance between the *C_α_* atoms of residue *i* of the J-domain and residue *j* of DnaK at time *t*. The dRMSs are then time-averaged over the last 10 ns of the MD trajectories (results reported in Tab. I).

**FIG. S1.**
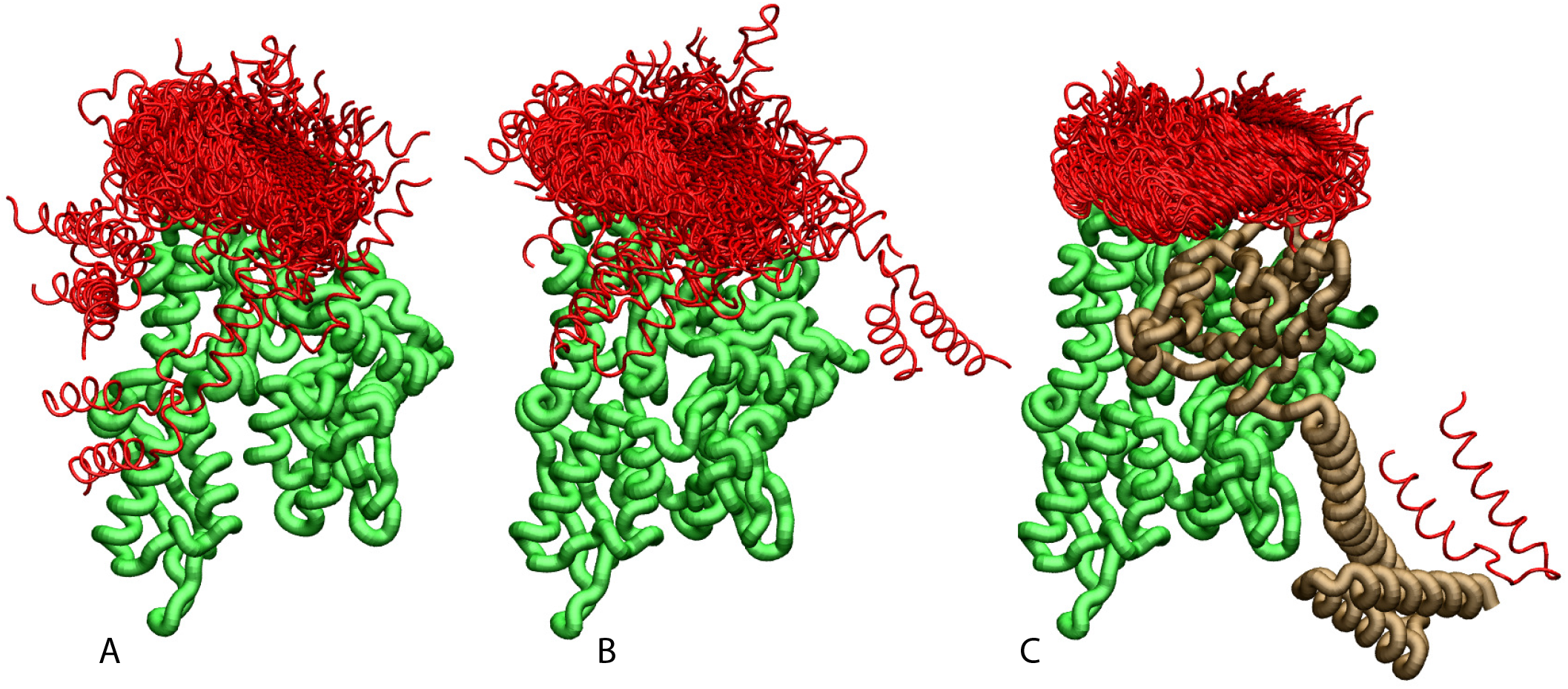
Ensemble of bound conformations predicted by coarse-grained simulations. The ensemble of bound JDs are reported for **A**) NBD(ADP), **B**) NBD(ATP) and **C**) FL(ATP). For ease of visualization, one out of 5 bound conformations is displayed. The JD is in red, the NBD in green and the SBD and linker in brown. All bound conformations have inter-protein distance of 8 Å or less and total binding energy below − 2*k_B_T*.

**FIG. S2.**
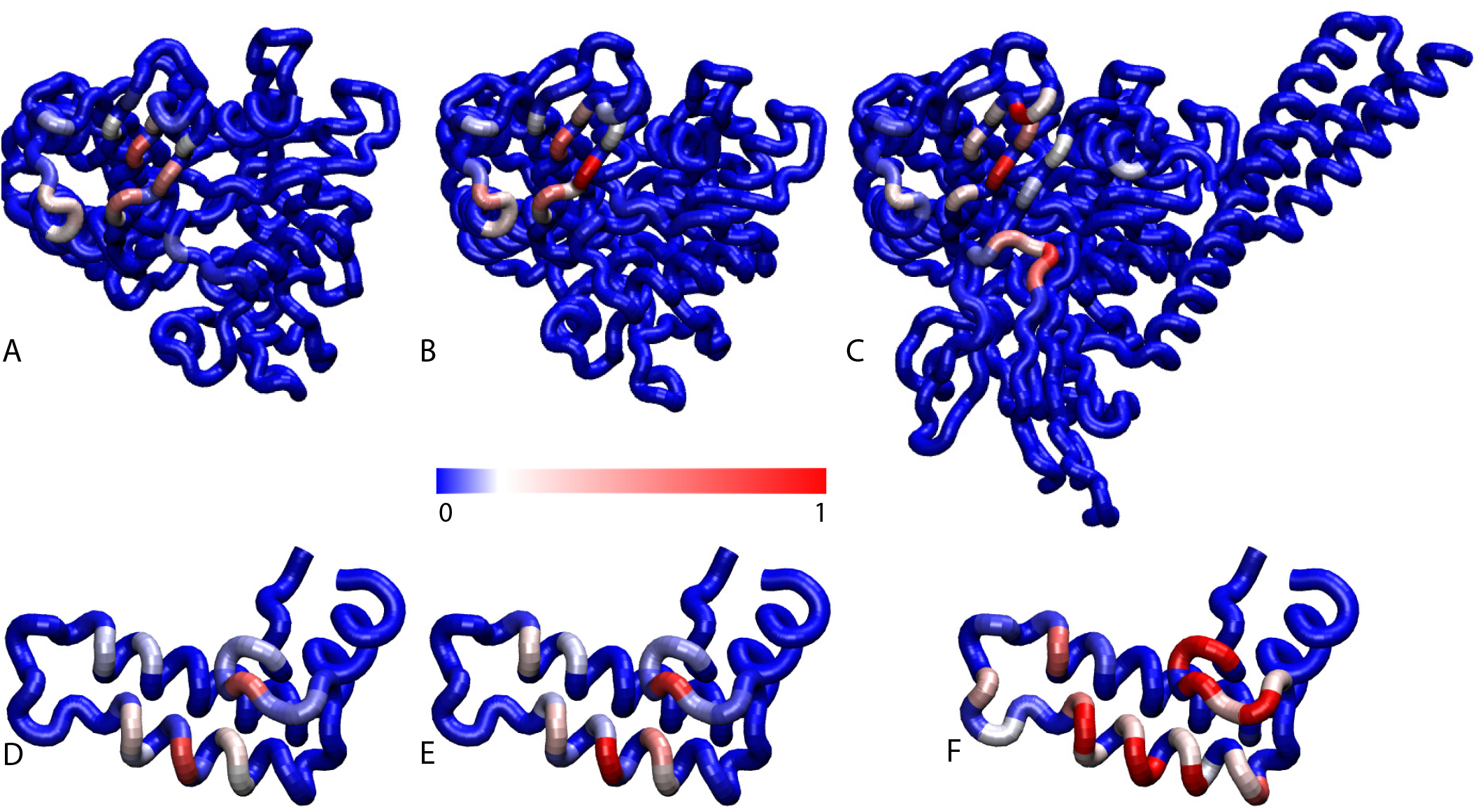
Contact frequencies predicted by coarse-grained simulations. Probabilities of DnaK residues to be in contact with the JD for **A**) NBD(ADP), **B**) NBD(ATP), **C**) FL(ATP). Probabilities of JD residues to be in contact with the DnaK for **D**) NBD(ADP), **E**) NBD(ATP), **F**) FL(ATP).

**FIG. S3.**
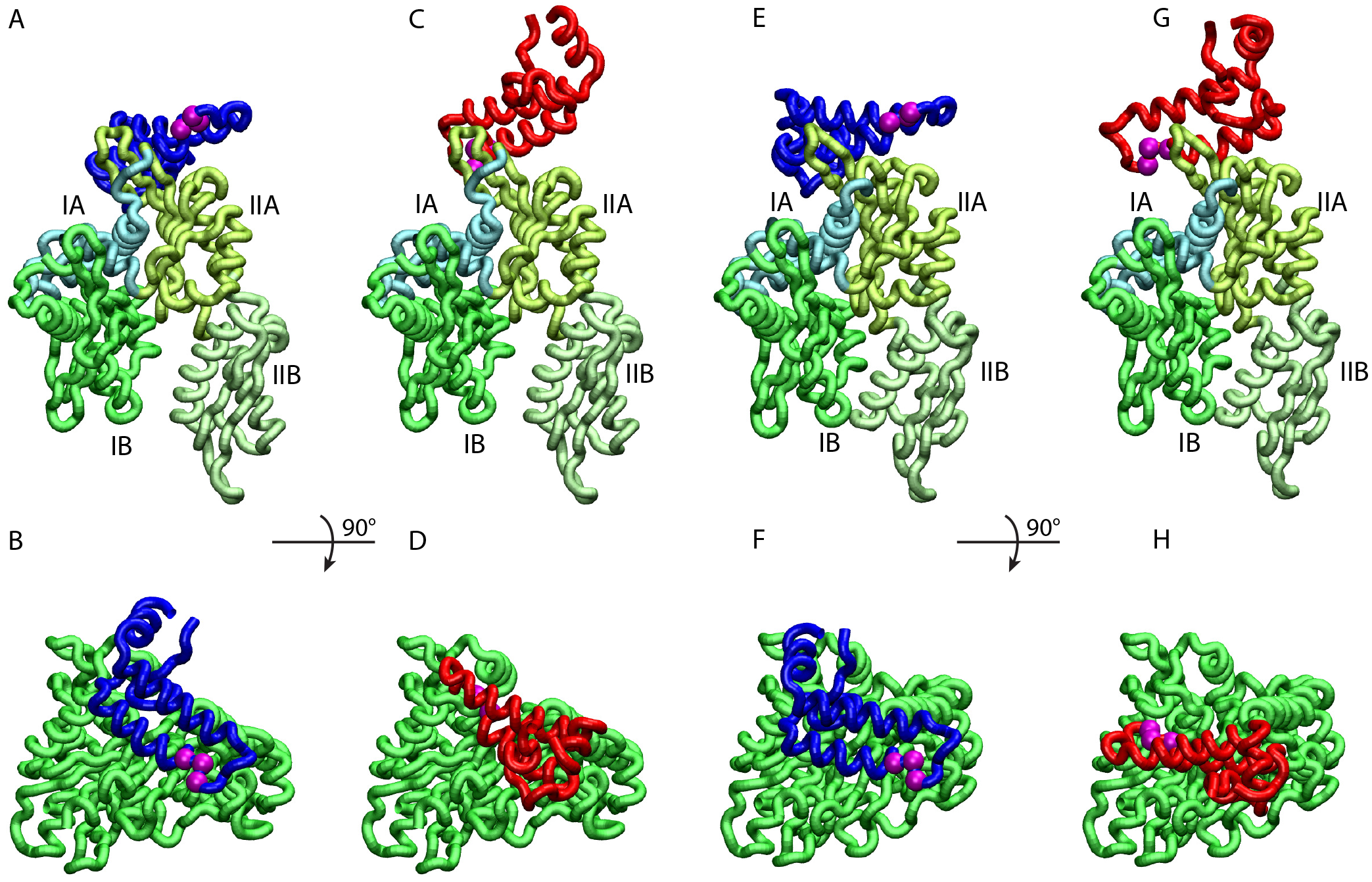
HPD-IN/OUT conformations for NBD(ADP) and NBD(ATP). Most representative HPD-OUT/IN conformations. **A,B**) HPD-OUT NBD(ADP), **C,D**) HPD-IN NBD(ADP), **E,F**) HPD-OUT NBD(ATP), **G,H**) HPD-IN NBD(ATP). The four lobes forming the sub-structures of the NBD are highlighted. The HPD tripeptide of the J-domain is depicted as magenta spheres. The JD is depicted in blue (HPD-OUT) or red (HPD-IN).

**FIG. S4.**
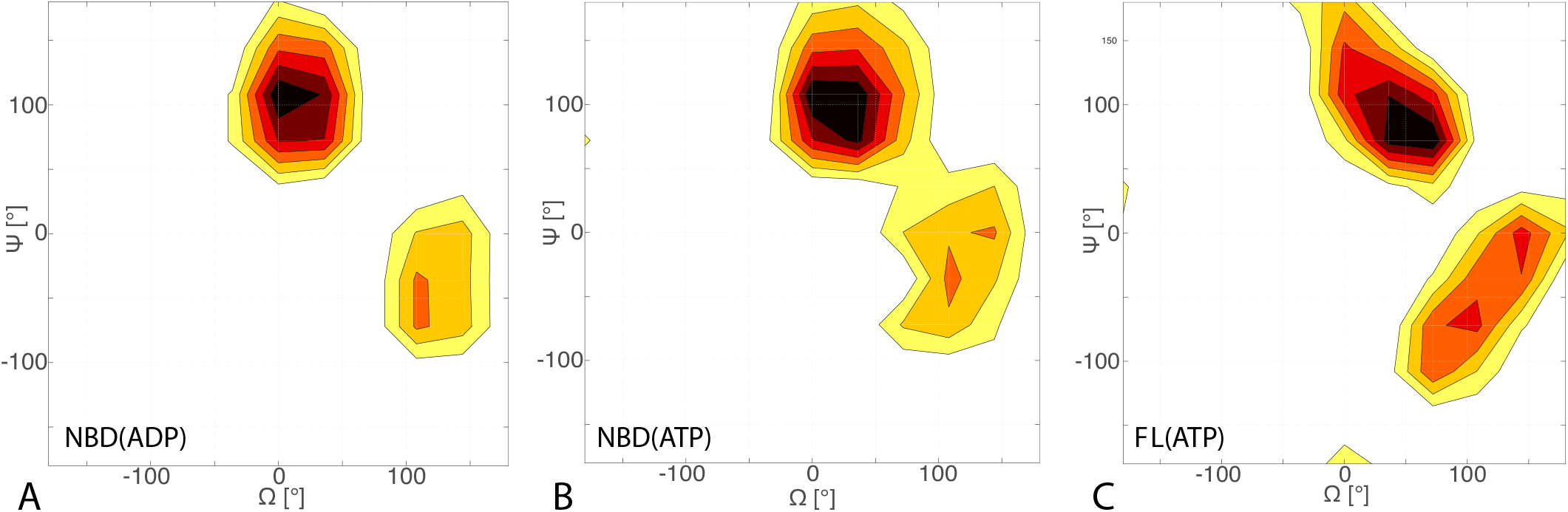
Free energy surface of bound CG conformations, using the third Euler angle. Free energy surface as a function of Euler angles (Ω, Ψ) defining the relative orientation of J-domain w.r.t. the DnaK NBD for **A**) NBD(ADP), **B**) NBD(ATP), **C**) FL(ATP). The fixed and rotating coordinate systems are defined by the inertia axes of the NBD and JD respectively. Iso-value lines are drawn at 1 (NBD(ADP/ATP)) or 1.2 (FL(ATP)) *k_B_T* intervals.

**FIG. S5.**
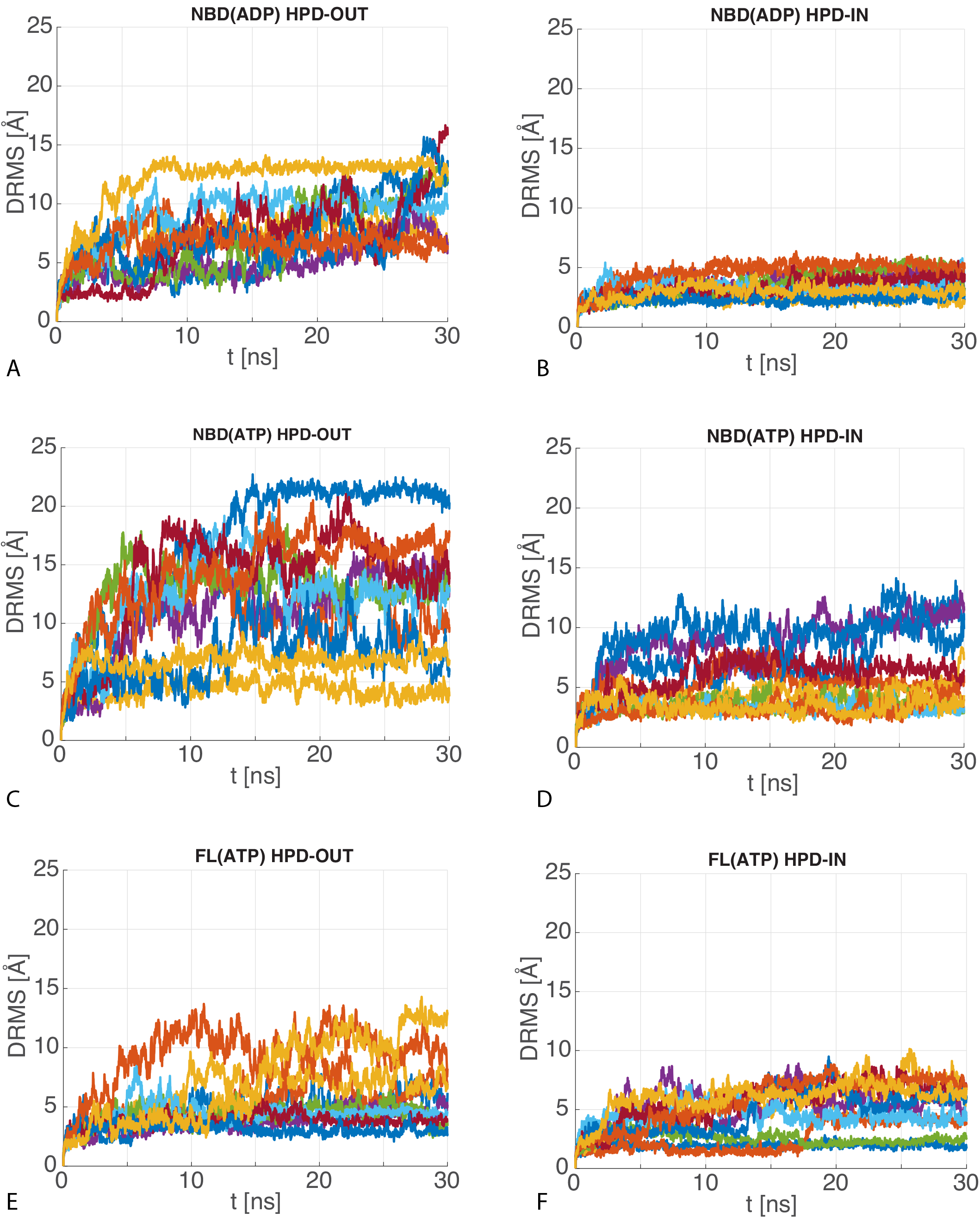
dRMS of Atomistic MD trajectories. The10 time series of the dRMS of the JD with respect to the initial JD configuration of each MD trajectory for **A**) HPD-OUT NBD(ADP), **B**) HPD-IN NBD(ADP), **C**) HPD-OUT NBD(ATP), **D**) HPD-IN NBD(ATP), **E**) HPD-OUT FL(ATP), **F**) HPD-IN FL(ATP).

**FIG. S6.**
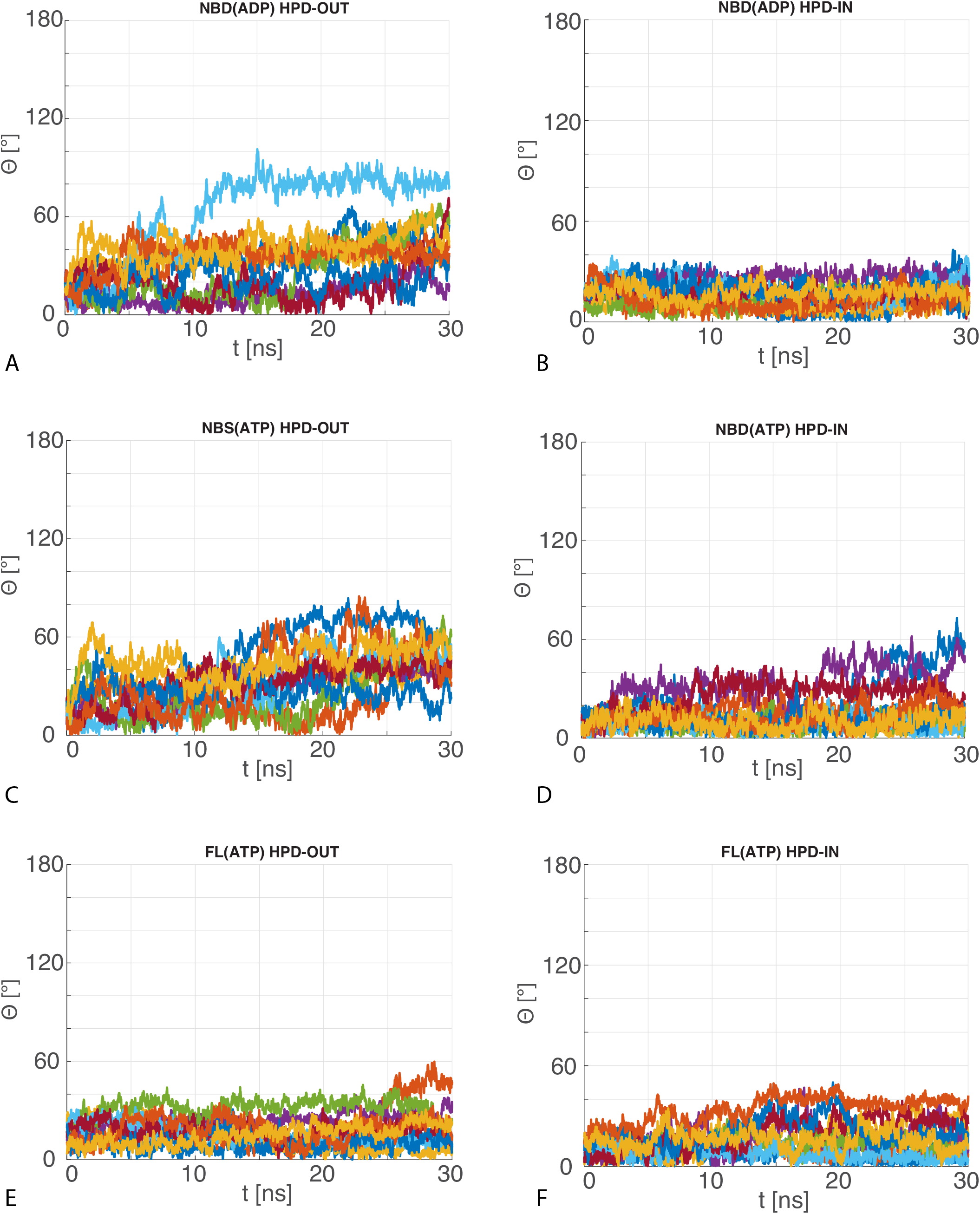
Angular deviation of Atomistic MD trajectories w.r.t. the central HPD-IN/OUT CG conformations. The 10 time series of the angle between the principal axis of the JD in the MD trajectories with respect to the principal axis of the most representative JD configuration in the CG ensemble for **A**) HPD-OUT NBD(ADP), **B**) HPD-IN NBD(ADP), **C**) HPD-OUT NBD(ATP), **D**) HPD-IN NBD(ATP), **E**) HPD-OUT FL(ATP), **f**) HPD-IN FL(ATP).

